# FetchR: an intuitive end-to-end solution for local RNA-Seq analyses

**DOI:** 10.64898/2026.07.30.741593

**Authors:** Dustin R. Fetch, Alexey A. Soshnev

## Abstract

RNA-Seq, analyses of RNA abundance by next-generation sequencing, has become a near-universal tool in modern biology. Availability of streamlined protocols and kits, straightforward ability to multiplex hundreds of samples, low cost of short-read sequencing, and well-established analytical pipelines make RNA-Seq a method of choice when even a few genes need to be analyzed in parallel. While many tools have been developed for quality control, mapping, and visualization of RNA-Seq data, managing all these individually still requires substantial familiarity with shell scripting and R, and remains a bottleneck for laboratories with limited computational background.

We assembled FetchR, an intuitive pipeline with built-in, clear explanations of features and outputs, for local analyses of RNA-Seq data from either own .fastq files or data imported from Sequence Read Archive via the ENA Portal API. The pipeline operates in Windows Subsystem for Linux (WSL) and is installed via a single script that handles all individual tools, as well as their dependencies and updates, including the reference genome annotation(s), and system requirements. The outputs include standard quality control checks, data visualization, read summation, differential gene expression analyses, visualization, and exploratory analyses using Gene Ontology and Gene Set Enrichment Analyses, as well as detailed logs of every step for subsequent reproducible reporting.

## 1. Introduction

Gene expression analyses is a key analytical tool to understand developmental transitions and mechanisms of disease (Bernadskaya and Christiaen, 2016). While the limitations of inferring cell identity from mRNA abundance are plenty, when costs, time, and required expertise are considered, no viable alternative comes close (Stark et al., 2019). As the throughput of short-read sequencing increased exponentially over the past decade, and many commercial providers offer turnaround times under a week for data delivery at prices competitive with the cost of reagents for a single quantitative PCR assay, analyses of RNA abundance by sequencing (RNA-Seq) will likely continue to dominate as first-line assay to understand the molecular phenotypes *in vivo*. Yet while the “wet-lab” portion of RNA-Seq has been streamlined to off-the-shelf kits, and sequencing process itself is typically outsourced to commercial service providers, analyses and data representation of an RNA-Seq experiment outcomes still represent a significant barrier.

Despite intuitive workflow and many excellent pipelines available, *e*.*g*. via Nextflow (Ewels et al., 2020), even deployment of well-documented tools requires the user to be comfortable with UNIX directory structure, command line, containers and version control – and in our anecdotal experience still represents a substantial barrier for users with limited experience in informatics. Alternative solutions include commercial software, often offered as a subscription service, or cloud-based tools which may offer limited analyses options, limited throughput, or be impractical if the original data must be analyzed locally due to privacy concerns.

FetchR lowers the barrier for entry into RNA-Seq analyses by the end user. While the pipeline provides clear step-by-step guidance to most steps and offers straightforward analyses options, it both allows for reasonably sophisticated exploratory analyses and data visualization, and offers extensive documentation for reproducible outputs for data presentation and reporting.

## 2. FetchR

Here, we briefly describe the pipeline requirements, capabilities and output, and demonstrate representative analyses using a publicly available RNA-Seq dataset.

### 2.1. System requirements and installation

FetchR is implemented in Windows Subsystem for Linux (WSL2), included by default in Windows 11. While WSL is also available for Windows 10 via winget, we envision that majority of potential FetchR users would be operating a Windows 11 machine. The system requirements are primarily driven by (a) size of unprocessed `.fastq` files dictating hard drive allocation, (b) preference for multithreaded processing, where parallelization improves overall speed in certain tasks, and (c) requirement for sufficient RAM imposed by STAR alignment software. Of note, FetchR automatically checks for available resources, informing decisions to allocate additional processor cores or memory to the process. While we recommend, at the minimum, HDD space exceeding the total `.fastq` file size at least four-fold, eight processor cores, and 32 Gb RAM for smooth operation of the resource-intensive part of the pipeline, the final steps encompassing all exploratory analyses can be run independently using a portable .`rds` file generated by the pipeline. In a real-life scenario, a single workstation-class PC can serve multiple investigators or projects for resource-intensive genome indexing and alignment part of the pipeline, and subsequent exploratory analyses can be completed on personal computers with minimal resource requirements.

FetchR installation is achieved through the execution of a single script, with several built-in checks. FetchR installer ensures that sufficient disk space is available (estimated at 8 Gb), and installation begins with necessary system wide packages (**Table 1**). With the machine prepared for general installation, Miniconda is installed, and a new `conda` environment, `RNAseqEnv`, is generated. All subsequent tools are installed within this environment (**Table 2**).

**Table 1:**
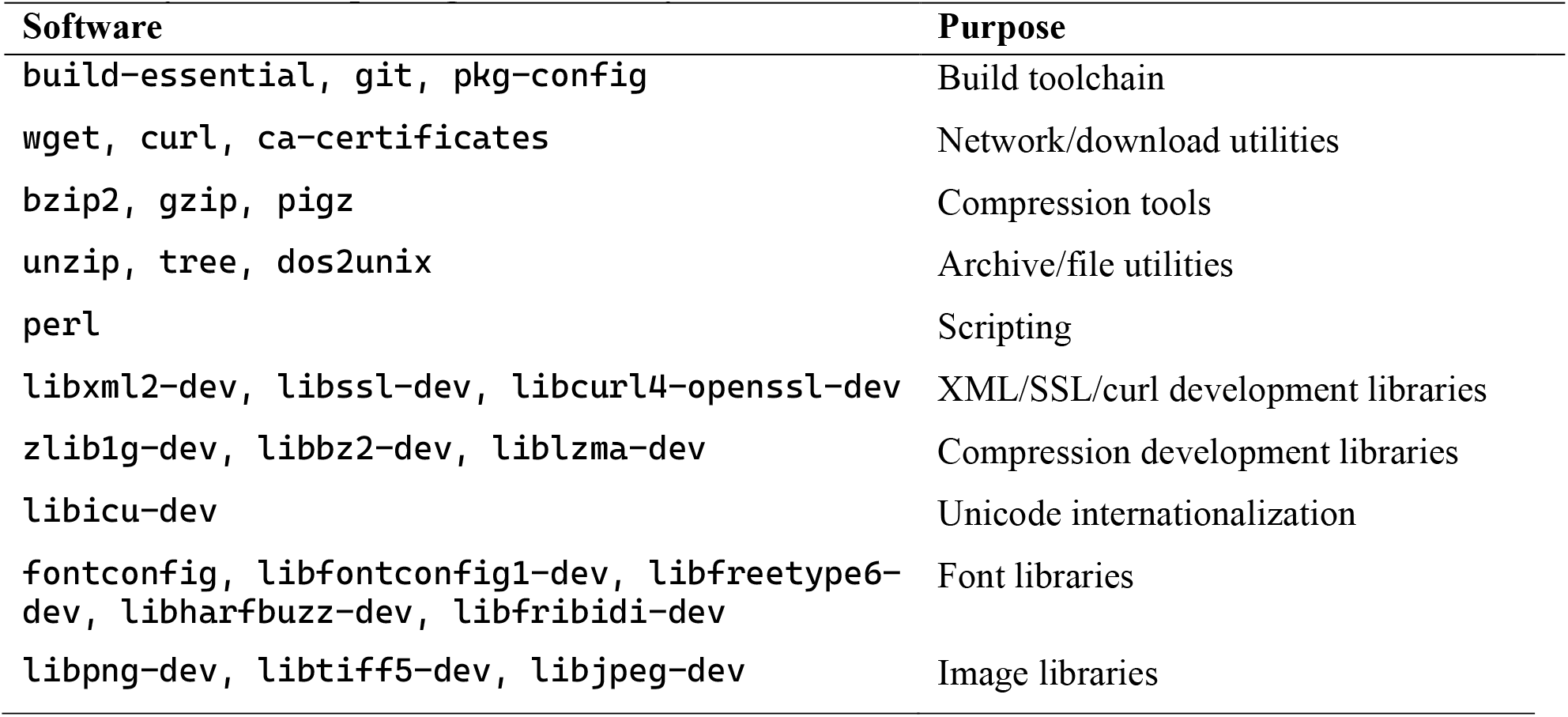
System-wide packages installed by FetchR.

**Table 2:** Tools installed within RNAseqEnv environment.

| Software | Current version | Purpose |
| --- | --- | --- |
| STAR | 2.7.11b | Alignment and genome indexing |
| subread | 2.1.1 | Provides featureCounts |
| samtools | 1.23.1 | BAM filtering, sorting, and indexing |
| fastqc | 0.12.1 | Pre-post trim quality control |
| multiqc | 1.35 | Aggregate QC reports |
| trim-galore | 0.6.11 | Reads trimmer |
| cutadapt | 5.2 | Trimming backend used by trim-galore |
| deeptools | 3.5.6 | Provides bamCoverage |
| Pigz, wget, curl, unzip, tree, dos2unix | unpinned | Make the resulting environment self-contained |
| r-base | 4.3 | R language runtime |
| pip | unpinned | Python package installer |

During the installation process, the pinned version of a given software is checked against current versions on `conda-forge` and `bioconda` channels (Gruning et al., 2018). If a newer version is available, the user is prompted with a choice to install either the known pinned version or the untested newer version. This installation stage ends with the installation of `r`, setting the stage for `r` packages installation (**Table 3**).

**Table 3.**
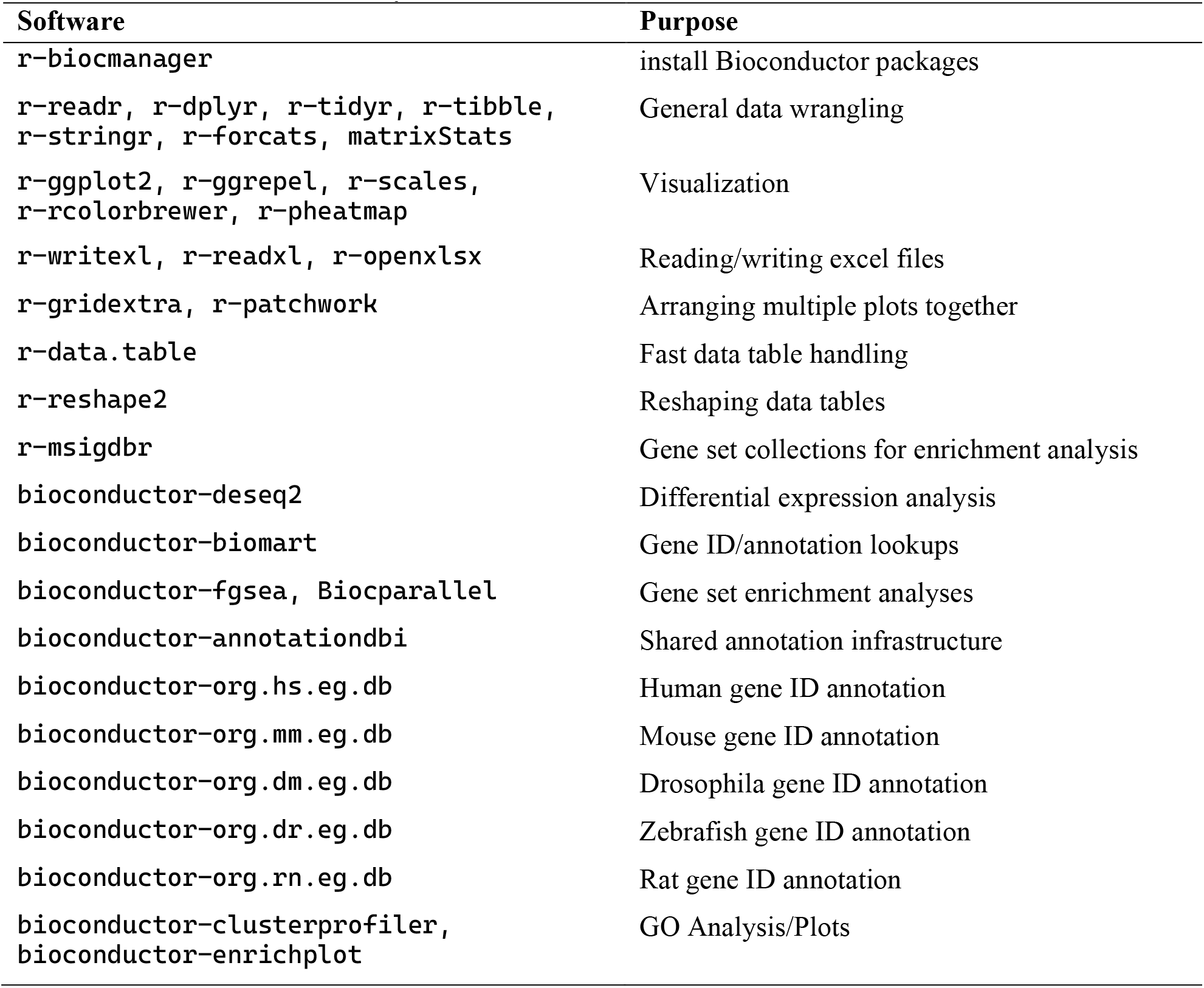
r packages installed by FetchR.

*Drosophila*, rat, and zebrafish gene ID annotations are considered optional by the installer, as such the failure of the environment to download the annotations will not result in a failed installation, and an attempt to retry a failed download of any. annotations will be made through `BiocManager` if this occurs. With all software installed, the installer writes a lightweight script, `rnaseq_check.sh`, which can be run to quickly verify the installation of all command line tools and `r` packages. A final verification of proper installation is confirmed, following which a full and minimal environment are saved, providing a useful starting point to clone the installation on other machines if desired. Importantly the installer script only installs what is necessary on the user’s machine, meaning that the installer will not reinstall already present software.

### 2.2. Data import and processing

With installation complete, FetchR uses a runner script, `FetchR_rnaseq_run.sh` for the processing of raw `.fastq` files into a `featureCounts` table ready for differential analysis (**Figure 1**). The runner script is compatible with `.fastq` files already present on the user’s machine as well as those accessible through an SRA study accession (SRP) or BioProject accession (PRJNA) which are retrieved using the ENA portal API (ref) and validated using md5 checksums provided by ENA. If accessing files through an SRP/PRJNA accession number, users are prompted to identify files of interest from the study as well as a location to download them to. If the files already exist on the machine, the user is asked to identify the folder where they are saved. FetchR automatically generates a project directory at a specified location for organized data export.

**Figure 1.**
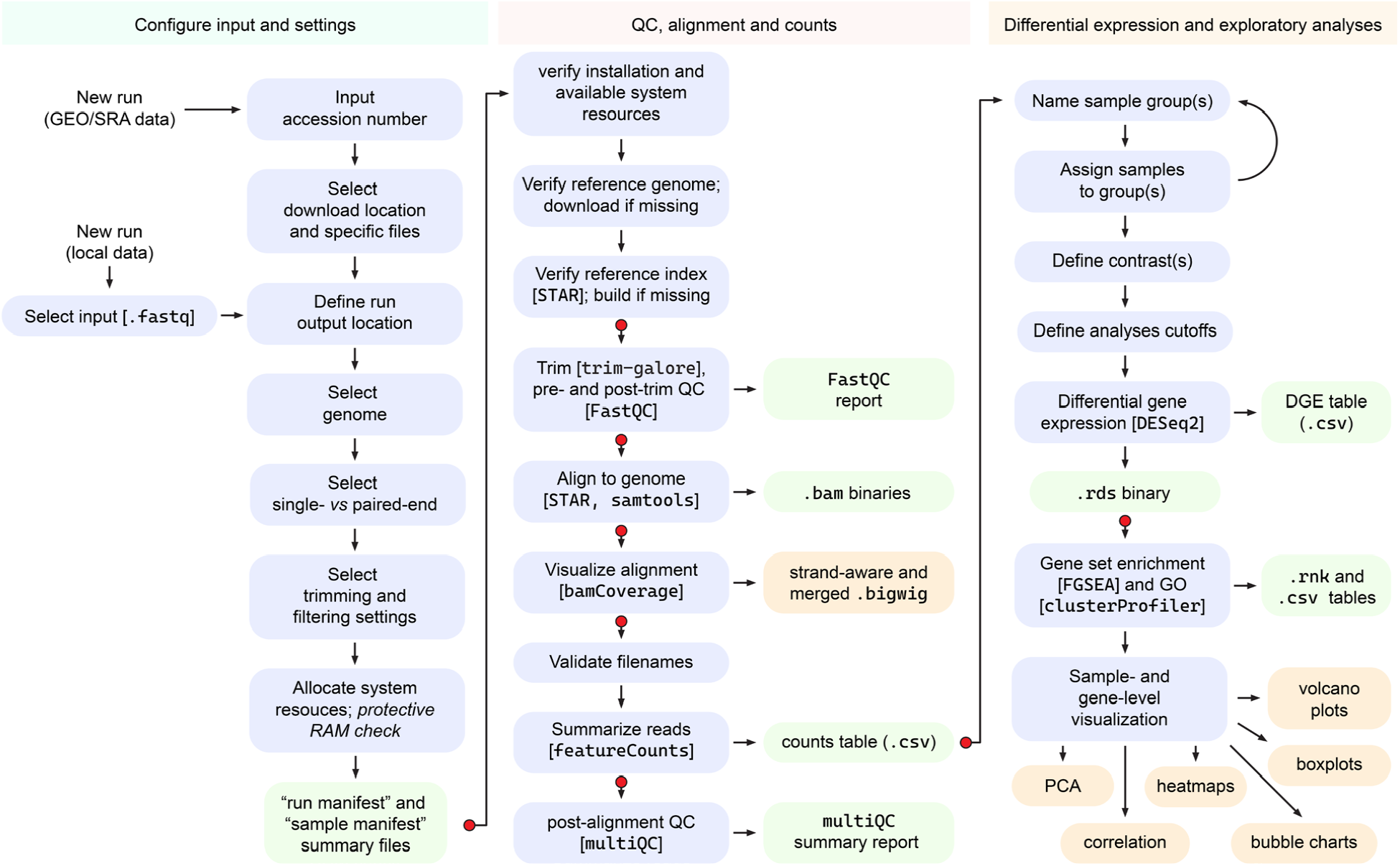
Workflow of FetchR. Input and settings are configured and saved in a “run manifest” and “sample manifest” files. Reference genome is installed and indexed if necessary, and .`fastq` files are processed with several consecutive QC checks, trimming, alignment and visualization via `FetchR_rnaseq_run.sh`. featureCounts output is analyzed via DESeq2 by `FetchR_rnaseq_diff_analysis.sh`, and exploratory analyses is completed via `FetchR_rnaseq_explore.sh`. Blue boxes indicate critical steps, green boxes denote significant outputs, and orange boxes show outputs that may be used in subsequent expert analyses. Red dots indicate select “resume points” where analyses can be restarted without re-doing previously completed steps.

With input and output locations specified, the user then chooses the genome that matches the samples, with current default options including hg38, mm10, rn7, dm6, danRer11, or “other genome”, the latter asking the user to provide a URL to download `.fasta` and .`gtf` annotation from. The user then specifies the download locations for genome(s), as well as where STAR index files will be generated (Dobin et al., 2013). Following this the user specifies their data’s read length, an appropriate STAR `sjdbOverhang` value, and whether the data is single- or paired-end. This information can automatically be propagated from SRA- or PRJNA-downloaded datasets. After providing a `featureCounts` strand setting (Liao et al., 2014), the user decides whether to run optional FastQC analysis pre- and/or post-trimming (Andrews, 2010), as well as any additional trimming and filtering settings (Krueger, 2026). With all settings determined, the user is presented with the total number of cores and threads available in their machine. Knowing this number, the user allocates threads to each processing step. As it is easy to use more RAM than their machine has available if executing multiple STAR jobs at once, FetchR compares a rough estimate of requested RAM required by STAR to what the machine has available. If necessary, FetchR will run in a low RAM mode, which trades off the run time for benefit of running on machines with as little as 32 Gb RAM. User then decides on whether to build `.bigwig` track files as well as the settings to do so (Ramirez et al., 2014). Finally, user can choose light, full, or no cleanup after the run is completed, removing none, some, or all of the intermediate files if the disk space is at a premium. The user is then presented with all discovered samples as well as an approval for the run overview.

At the start of every run, a “run manifest” and “sample manifest” are generated as a record of run settings and the samples that were processed using these settings. The runner script then checks available disk space as well as all necessary software is present. Reference genome `.fasta` and `.gtf` files are confirmed, or if missing, downloaded. Similarly STAR indexed genomes, with the proper overhang, are confirmed at the specified location, and, if not found, are built automatically. Finally, expected `.fastq` files are verified and the processing run begins. A run can be cancelled and resumed at any point using the “resume option” present in the runner, which utilizes the resume state file written at the start of every run. The user then chooses one of nine predefined stages to restart processing from, with an opportunity to reallocate computer resources if desired.

### 2.3. Differential gene expression

The `featureCounts` output table is then piped into the differential analysis script, `FetchR_rnaseq_diff_analysis.sh`. Initialization of this script requires the user to identify the location of an output `featureCounts` file, followed by organism selection. The user then assigns samples to groups which are used to define pair-wise comparisons of interest. The user can set as many contrasts as needde, indicating which group represents the contrast and control for each comparison. FetchR allows for adjustable comparison values including false discovery rate cutoff, log fold change cutoff, and FGSEA/GO specific settings. After run initialization, a log file is written which contains the `r` version, `r` package versions, OrgDb data vintage, and all user-set parameters.

After run information is archived the raw `featureCounts` table is stripped down, passing the counts matrix and sample metadata to DESeq2 (Love et al., 2014), which performs size factor estimation as a form of normalization, and following that fits the data with a negative binomial generalized linear model, and estimates dispersion for each geneThe script then connects to OrgDb package to map counts to Entrez IDs. Importantly, Entrez IDs that are identified here become the GO background universe, an optional but recommended set of genes used for gene ontology analyses. DESeq2 fits a global model of all samples which is used for VST transformation and saved within the later created “explorer bundle”. From this global model all comparisons are extracted.

The Wald test statistic is saved to a `.rnk` file, gene symbols are mapped onto the ranked list, and then are immediately piped into FGSEA (Korotkevich et al., 2021) which runs the pre-ranked data against seven collections; Hallmark, GO Biological Processes, GO Cellular Component, GO Molecular Function, Transcription Factor and microRNA targets, Oncogenic Signatures, and Cell Type Signatures (Liberzon et al., 2015). GO enrichment (ORA) is similarly run, providing a complementary approach to generated FGSEA data. Heatmaps, volcano plots, and Spearman rank correlations are calculated and provided for every contrast. This script finalizes with the saving of an explorer `.rds` file, which allows for exploration of the data in greater detail in low-resource environment (*e*.*g*. school laptop).

### 2.4. Exploratory analyses

The explorer script allows the user to perform expert-level analyses in an interactive fashion, investigating either sample-level patterns, or focusing on genes of interest. Information is displayed via several standard graphs that cnja be customized top specific needs (**Figure 2**). This script is reliant on the `explorer_bundle.rds` file generated at the end of the `diff_analysis` script. This bundle is a globally fitted DESeq2 object that contains the full-dataset VST matrix, sample metadata, group/contrast information, and gene annotation mappings. Importantly the explorer bundle prevents users from repeating read processing, DESeq2 model fitting, or VST calculation. PCA plots, expression heatmaps, and sample-correlation heatmaps are generated using this previously calculated VST matrix. Gene expression boxplots are generated from DESeq2-normalized counts estimated from the original global model fit. The explorer uses data from the explorer bundle to perform GO enrichment, pre-ranked GSEA, as well as customization of all above-described plots. The purpose of this script is customization of the basic plots expected or wanted from a bulk RNA-Seq experiment. This includes the highlighting of genes of interest, ability to explore and change LFC/FDR values, and to explore sample-level relationships. As the `.rds` file is portable, and the script is relatively lightweight, this provides the end users with the ability to explore their data independent of workstation machines.

**Figure 2.**
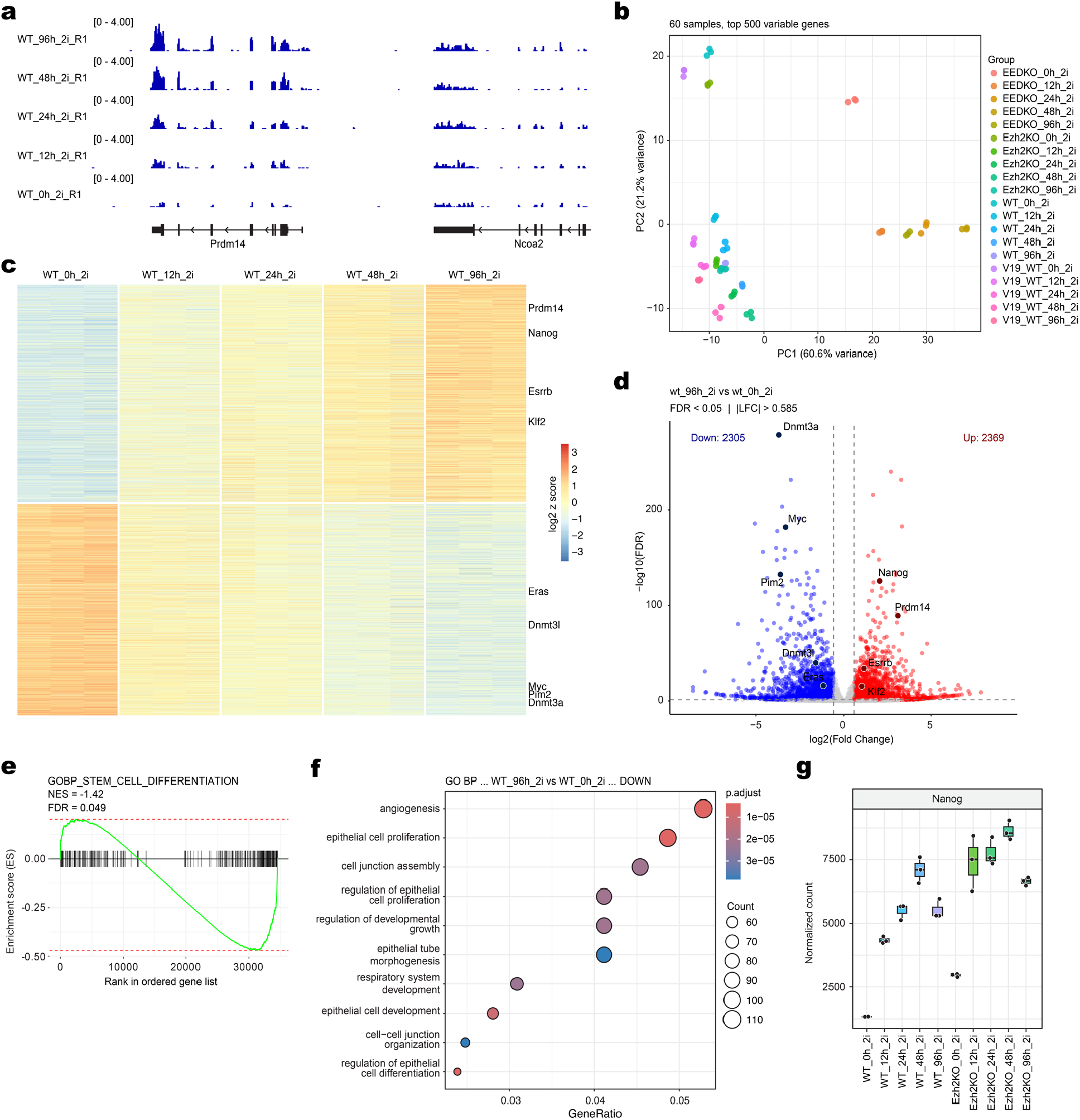
Outputs of FetchR. **a**, .bigwig visualization of individual replicates; **b**, principal component analyses plot for all samples in the dataset; **c**, heatmap of differentially expressed genes within contrast of interest, with select genes indicated; **d**, volcano plot for user-defined contrasts; **e**, GSEA analyses output; **f**, gene ontology bubble plot; **g**, box plots of normalized counts of a single gene of interest across several conditions and genotypes. All examples shown here represent direct output of FetchR minimally adjusted in Adobe Illustrator, using RNA-Seq data from (Chapa et al., 2025), downloaded from Gene Expression Omnibus GSE143293.

## 3. Concluding remarks

FetchR was designed to make RNA-Seq analyses accessible to a user with limited experience in bioinformatics. While it is not intended to substitute the knowledge of statistics and informatics, we envision it will bridge the gap and provide a first glimpse at the possibilities of genome-wide analyses for a new user. Like the commercial wetlab kits are not intended to substitute the knowledge of molecular biology and biochemistry, yet provide a helpful time saver in wet-lab experiments, we hope this software will expedite and streamline RNA-Seq analyses for some end users.

## Acknowledgements

We thank the members of the Soshnev and Barton laboratories at UT San Antonio for helpful suggestions and testing of FetchR. The code was written with the assistance of Claude AI. The authors take full responsibility for the code and its annotation.

## Author contributions

D.R.F. conceived the project and designed FetchR with input from A.A.S; D.R.F and A.A.S jointly wrote the manuscript.

## Data and code availability

FetchR is available at https://github.com/DustinFetch/FetchR-RNAseq-Pipeline.git

